# Tracking *Borrelia afzelii* from infected *Ixodes ricinus* nymphs to mice suggests a direct ‘gut-to-mouth’ route of Lyme disease transmission

**DOI:** 10.1101/316927

**Authors:** Tereza Pospisilova, Veronika Urbanova, Ondrej Hes, Petr Kopacek, Ondrej Hajdusek, Radek Sima

## Abstract

Quantitative tracking of *Borrelia afzelii* has shown that its transmission cycle differs from the salivary route of *B. burgdorferi* transmission by *Ixodes scapularis*. *Borrelia afzelii* are abundant in the guts of unfed *Ixodes ricinus* nymphs and their numbers continuously decrease during feeding. In contrast, spirochetes are not present in the salivary glands. *Borrelia afzelii* transmission starts during the early stages of feeding, spirochetes could be detected in murine skin within 1 day of tick attachment. Tick saliva is not essential for *B. afzelii* infectivity, the main requirement for successful host colonization being a change in outer surface protein expression that occurs in the tick gut during feeding. Spirochetes in vertebrate mode are able to survive within the host even if the tick is not present. On the basis of our data we propose that a direct ‘gut-to-mouth’ route of infection appears to be the main route of *B. afzelii* transmission.

**Importance:** Lyme borreliosis is the most common vector-borne disease in the USA and Europe. The disease is caused by the *Borrelia* spirochetes and is transmitted through *Ixodes* ticks. A better understanding of how *Borrelia* spirochetes are transmitted is crucial for development of efficient vaccines for preventing Lyme borreliosis. Here we present that the transmission of European *B. afzelii* spirochetes by *I. ricinus* ticks significantly differs from the model transmission cycle described for American *B. burgdorferi*/*I. scapularis*. We suggest that *B. afzelii* is not transmitted via salivary glands but most likely through the ‘midgut to mouthpart’ route. We further demonstrate that tick saliva is not important for *B. afzelii* transmission and infectivity. Therefore, we support early studies by Willy Burgdorfer, who proposed that *Borrelia* transmission occurs by regurgitation of infected gut contents. Our findings collectively point to the *Borrelia*-tick midgut interface as the correct target in our endeavours to combat Lyme borreliosis.

## Introduction

Lyme borreliosis is the most common vector-borne disease in Europe and the USA. It is caused by the spirochetes *Borrelia burgdorferi* in the USA or by the *B. burgdorferi* sensu lato (s.l.) complex comprising *B. afzelii, B. garinii*, and *B. burgdorferi* in Europe. *Borrelia* spirochetes are maintained in nature through an enzootic cycle involving small vertebrates, primarily rodents and birds, and are vectored by ticks of the genus *Ixodes* (1).

Understanding the complex interactions within the tick-*Borrelia*-host triangle is indispensable for the development of efficient vaccines or drugs against Lyme disease. Progress in understanding borreliosis transmission has been achieved during the last three decades, mainly in the USA, by investigation of *B. burgdorferi* strains vectored by *I. scapularis*. Three possible hypotheses for *Borrelia* transmission were proposed in the early studies: (i) the first observations favored a direct infection via mouth parts by regurgitation of the spirochetes present in the midgut contents (2); (ii) a salivary route of transmission that assumed systemic distribution of spirochetes within the tick body (3); (iii) infection via contaminated faeces was also considered (2, 4) but soon abandoned (5). A number of following studies corroborated the salivary route of *B. burgdorferi* transmission by *I. scapularis* as the most likely. Therefore, the current, generally accepted model of Lyme disease transmission can be summarized as follows: At the beginning, larval *I. scapularis* acquire *Borrelia* spirochetes from infected vertebrate reservoir hosts. *Borrelia burgdorferi* spirochetes then multiply rapidly in feeding larvae and during the first days post-repletion. The number of spirochetes are then dramatically reduced during subsequent molting (6). Spirochetes persisting in the nymphal midgut upregulate OspA (7), and stay attached to the TROSPA receptor on the surface of the midgut epithelial cells (8). Spirochetes remain in this intimate relationship until the next blood meal. As the infected nymphs start feeding on the second host, *Borrelia* spirochetes sense appropriate physiochemical stimuli that trigger their replication (7, 9). Their numbers increase exponentially (10, 11), and the spirochetes downregulate OspA and upregulate OspC (7, 12). Simultaneously, ticks downregulate the production of TROSPA (8). These changes help spirochetes to detach from the midgut, penetrate into the hemolymph, migrate to the salivary glands (8) and infect the vertebrate host.

Understanding of Lyme borreliosis in Europe lags far behind the USA, mainly because the situation is complicated by the existence of several different species in the *B. burgdorferi* s.l. complex that act as causative agents of the disease. To date, only a few papers have been published regarding transmission of *B. burgdorferi* s.l. strains by *I. ricinus* ticks, suggesting that the transmission of European *Borrelia* strains might differ from the model cycle described for *B. burgdorferi*/*I. scapularis* (see (5) for review).

In this study, we have performed a quantitative tracking of *B. afzelii* from infected mice to *I. ricinus* and back to naïve mice. We further tested the role of tick saliva in infectivity and survival of *B. afzelii* spirochetes. Based on our data, we proposed the concept of a direct ‘gut-to-mouth’ route of *B. afzelii* transmission from infected *I. ricinus* nymphs to the host.

## Results

### *Borrelia afzelii* – *Ixodes ricinus* transmission model

In order to understand the Lyme disease problem in Europe, the development of a transmission model is essential for the European vector *I. ricinus* and local *Borrelia* strains of the *B. burgdorferi* s.l. complex. For this purpose, we established a novel, reliable and robust transmission model employing C3H/HeN mice, *I. ricinus* ticks and the *B. afzelii* CB43 strain isolated from local ticks (13). This strain develops systemic infections in mice and causes pathological changes in target tissues. Variably intensive lymphocytic infiltrations were detected in the heart, where the majority of inflammatory cells were concentrated in the subepicardial space with infiltration of myocytes (Fig. 1A). Inflammatory infiltration was prominent within the urinary bladder. The most prominent changes were in the submucosa, close to the basal membrane (Fig. 1B). In the skin, weak infiltration of the epidermis and dermis was documented, however most lymphocytes were found in deep soft tissues (Fig. 1C). *Borrelia afzelii* CB43 also turned out to be highly infectious for *I. ricinus* ticks as positive infection was detected in 90-100% of molted nymphs that fed on infected mice as larvae.

**FIG 1.**
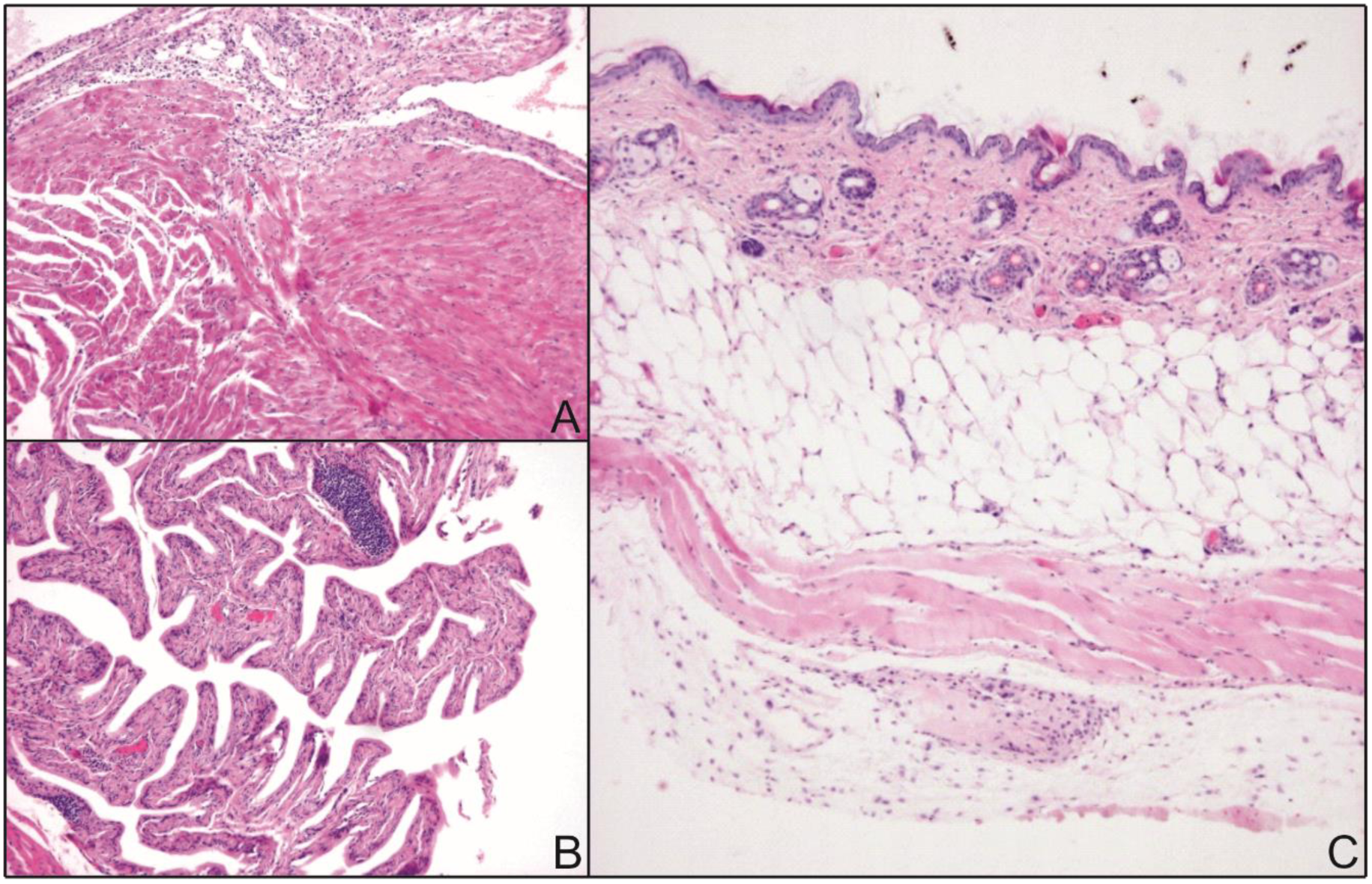
Pathological changes in target tissues of mice with *B. afzelii* infection. (**A**) Low power section shows lymphocytic infiltration in the subepicardial space. (**B**) Urinary bladder mucosa shows lymphoid infiltration, including dense lymphoid aggregates within the submucosa. (**C**) Lymphocytic infiltrates are not prominent in the skin, and the majority of lymphoid infiltration is located deep in the connective tissue.

### *B. afzelii* population grows rapidly in engorged *I. ricinus* larvae and during molting to nymphs

Studies on the dynamic relationship between the Lyme disease spirochete and its tick vector were previously performed on an *I. scapularis*/*B. burgdorferi* model (6, 10). Nevertheless, little is known about the growth kinetics of European *B. afzelii* in *I. ricinus* ticks. The number of spirochetes was determined in engorged *I. ricinus* larvae fed on *B. afzelii*-infected mice and then at weekly intervals until larvae molted to nymphs. Measurements were completed at the 20th week post-molt. The mean number of spirochetes in fully fed *I. ricinus* larvae examined immediately after repletion was relatively low, 618 ± 158 (SEM) spirochetes per tick. Then the spirochetes multiplied rapidly in engorged larvae and their numbers continued to increase during molting to nymphs. The maximum number of spirochetes, 21 005 ± 4 805 (SEM) per tick, was detected in nymphs in the 2nd week after molting. Spirochetal proliferation then halted and the average spirochete number became relatively stable from the 4th to 20th weeks post-molt, slightly oscillating around the average number of about 10 000 spirochetes per tick (Fig. 2).

**FIG 2.**
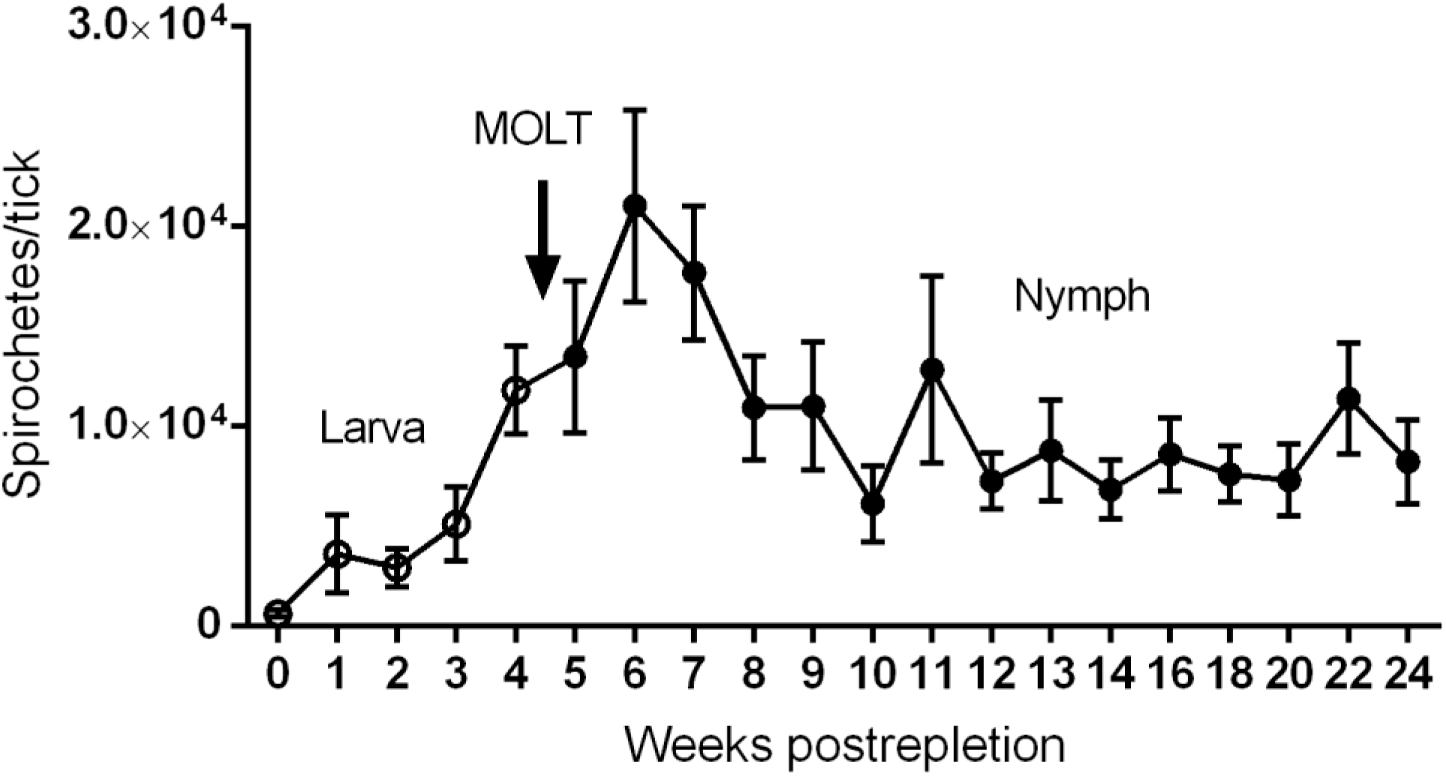
*B. afzelii* growth kinetics during larval-to-nymphal development in *I. ricinus*. Spirochetes multiply in engorged larvae, as well as during molting to nymphs. Then, spirochetal proliferation stops and spirochete numbers stay relatively stable from the 4th to 20th week post-molt. Each datapoint represents a mean of 20 individually analyzed ticks, and bars indicate standard errors of means.

### *B. afzelii* numbers in *I. ricinus* nymphs dramatically drop during feeding

We further examined the absolute numbers of *B. afzelii* spirochetes in infected *I. ricinus* nymphs during feeding. Nymphs were fed on mice, forcibly removed at time intervals 24, 48 and 72 hours after attachment and the spirochetes were then quantified by qPCR. Prior to feeding, the mean number of spirochetes per nymph was 10 907 ± 2 590 (SEM). After 24 hours of the tick feeding, the number of spirochetes decreased to 7 492 ± 3 294 (SEM). In the following 2nd and 3rd day of blood intake, the numbers continued to drop to 2 447 ± 801 (SEM) and 720 ± 138 (SEM) spirochetes per tick, respectively (Fig. 3). As this result was in striking contrast to the reported progressive proliferation of *B. burgdorferi* during *I. scapularis* nymphal feeding (10, 11), we confirmed the gradual decrease in *B. afzelii* spirochetes in the midguts of feeding *I. ricinus* nymphs using confocal immuno-fluorescence microscopy. By contrast, a parallel examination of the salivary glands from the same nymphs demonstrated that no spirochetes were present in this tissue at any stage of feeding (Fig. 4).

**FIG 3.**
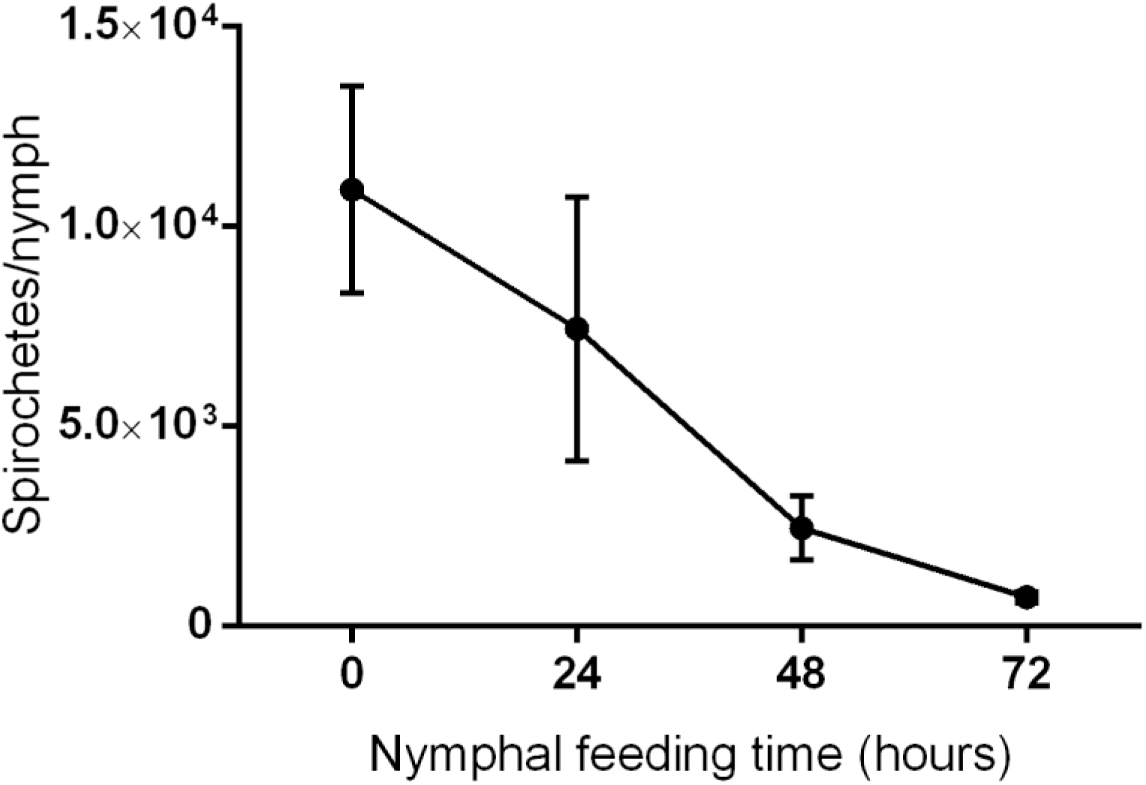
*B. afzelii* kinetics during nymphal feeding. During feeding, spirochete numbers continuously decrease from 10^4^ spirochetes/tick at the beginning, to several hundred spirochetes/tick at the end of nymphal feeding. Each datapoint represents the mean of 20 individually analyzed ticks, and bars indicate standard errors of means.

**FIG 4.**
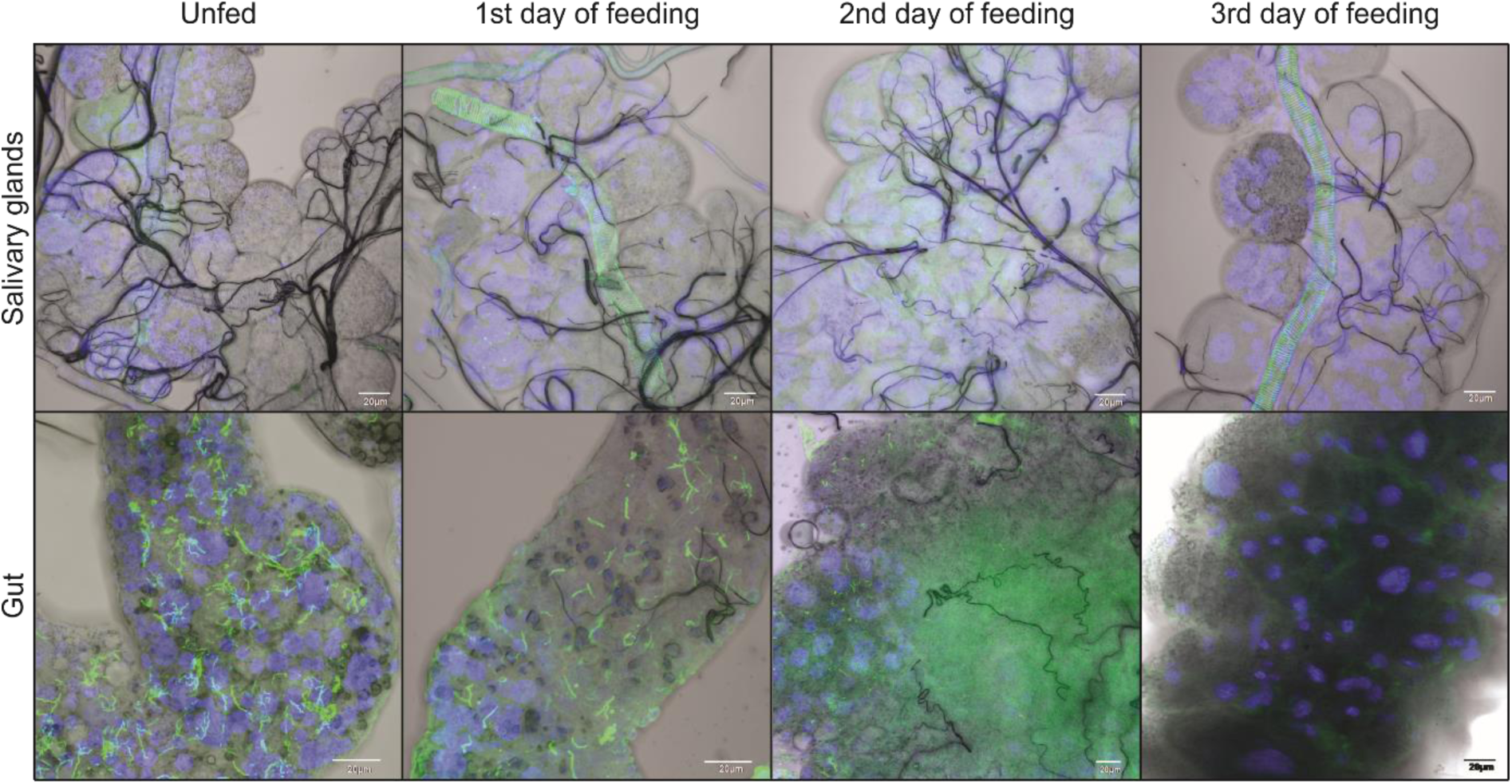
Presence of *B. afzelii* spirochetes in guts and salivary glands of feeding *I. ricinus* nymph. Spirochetes are clearly visible in midguts of *B. afzelii* infected nymphs. Their numbers significantly decrease during feeding. In contrast, spirochetes are hardly detectable in salivary glands of feeding *I. ricinus* nymph. *B. afzelii* spirochetes are stained with anti-borrelia antibody (green); nuclei are stained with DAPI (blue).

### Ability of *B. afzelii* spirochetes to develop a chronic infection in mice increases with feeding time

It is generally known that the risk of acquiring Lyme disease increases with the length of tick feeding (5). In subsequent experiments, we focused on the infectivity of *B. afzelii* transmitted via *I. ricinus* nymphs. To determine the minimum length of tick attachment time required to establish a permanent infection in mice, *B. afzelii* infected nymphs were allowed to feed on mice for 24, 48, and 72 hours (10 nymphs per mouse). The ability of *B. afzelii* spirochetes to promote a chronic infection increased with the length of tick attachment. All mice exposed to the bite of *B. afzelii* infected ticks for 24 hours remained uninfected, whereas 8/10 mice exposed for 48 hours and 10/10 mice exposed for 72 hours became infected. These results show that the time interval between 24 and 48 hours of exposure to the *B. afzelii* infected tick is critical for the development of a systemic murine infection.

### *B. afzelii* spirochetes are already present in the murine dermis on the first day of tick feeding

The delay in development of a *B. afzelii* infection in mice may support the notion that the spirochetes are still “on the road” towards the tick salivary glands during the first day after attachment. To test this hypothesis, we determined the number of *B. afzelii* in murine skin biopsies from the tick feeding site at time intervals of 24, 48 and 72 hours after feeding. Surprisingly, skin biopsies from 9/10, 10/10, and 10/10 mice were PCR positive at time intervals of 24, 48 and 72 hours, respectively. Analysis by qPCR further revealed that there were no significant differences in the number of spirochetes in skin samples at defined time intervals (Fig. 5A). This intriguing result was also confirmed by confocal microscopy, revealing clearly the presence of spirochetes in murine skin biopsies during the first day of tick feeding (Fig. 5B). Together with the rapid decrease in spirochetal number in nymphal midguts during feeding (Figs. 3 and 4), these results imply that the massive migration of spirochetes to the host commences soon after the blood meal uptake.

**FIG 5.**
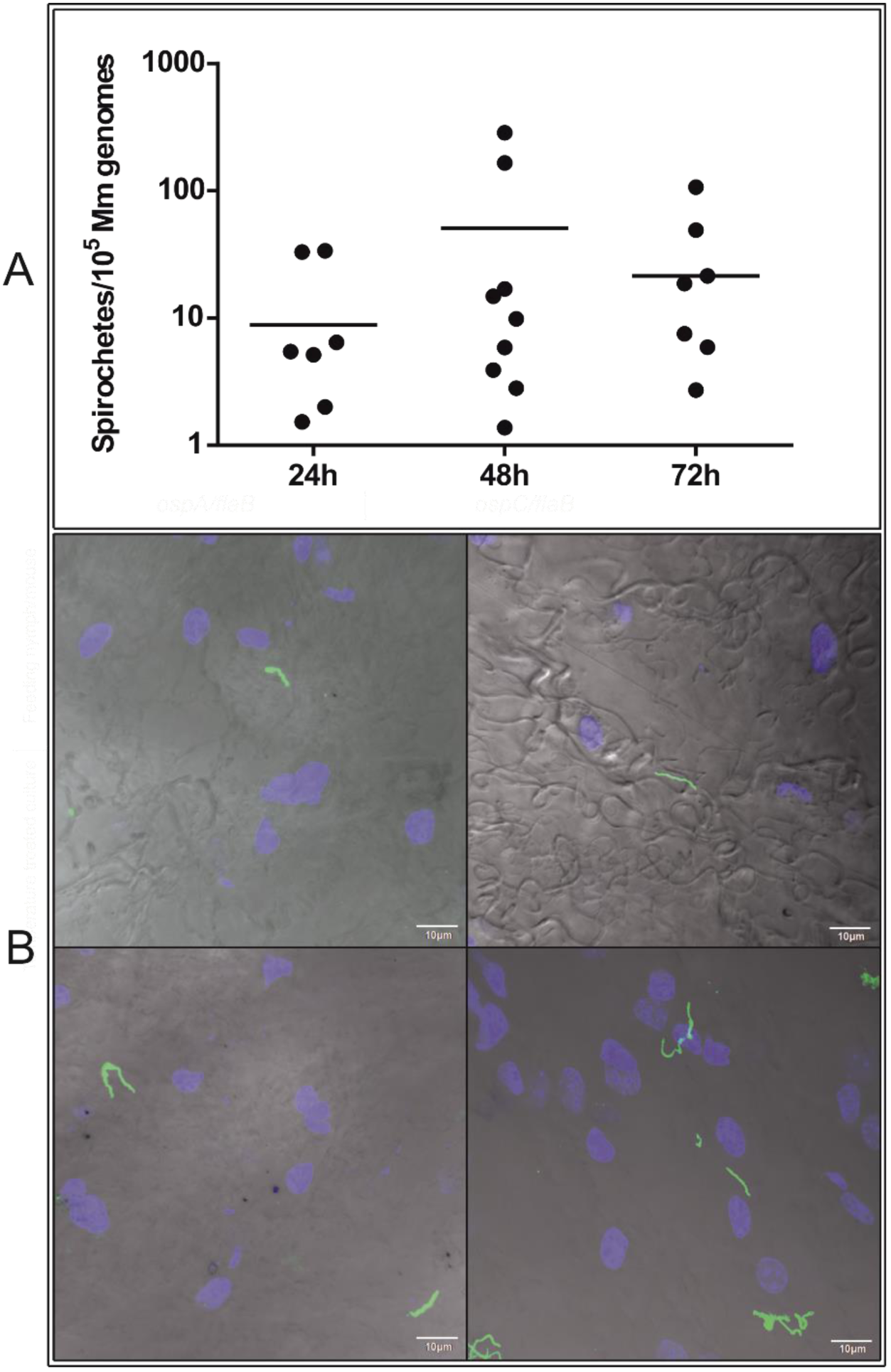
Timing of *B. afzelii* transmission from *I. ricinus* nymph to mouse. Skin biopsies from mice exposed to infected ticks for various time periods were tested for infection by qPCR (**A**) or confocal microscopy (**B**). *B. afzelii* spirochetes are present in the skin at the early stages of tick feeding. (**A**) Each datapoint represents the number of *B. afzelii* spirochetes/10^5^ murine genomes in individually analyzed skin biopsies. (**B**) Presence of *B. afzelii* spirochetes in murine skin at the 24 hour time point. *B. afzelii* spirochetes are stained with anti-*Borrelia* antibody (green); nuclei are stained with DAPI (blue).

### Tick saliva does not protect the early *B. afzelii* spirochetes against host immunity

The apparent contradiction between the early entry of *B. afzelii* spirochetes into the vertebrate host and their delayed capability to develop a permanent infection supports the concept of the tick saliva role in the successful dissemination and survival of spirochetes within the host body. In order to verify that tick saliva is essential for *B. afzelii* survival in the mice, we designed and performed the following experiment. In experimental group 1, uninfected *I. ricinus* nymphs (white labelled) were allowed to feed simultaneously with *B. afzelii*-infected nymphs (red labelled) at the same feeding site. After 24 hours of cofeeding, *B. afzelii*-infected nymphs were removed, while uninfected ticks fed on mice until repletion and served as a source of saliva. In control group 1, *B. afzelii*-infected nymphs fed for 24 hours without any support of uninfected ticks. In control group 2, *B. afzelii*-infected nymphs were allowed to feed until repletion. Four weeks later, *B. afzelii* infections in ear, heart and urinary bladder biopsies were examined by PCR. No infection was detected in any of examined tissues in experimental and control group 1, where the infected ticks fed for only 24 hours. By contrast, all tissues were PCR positive in control group 2, where the infected nymphs fed until repletion (Fig. 6A). These results revealed that the presence of uninfected ticks and their saliva is not sufficient to protect early spirochetes against elimination by the host immune system.

**FIG 6.**
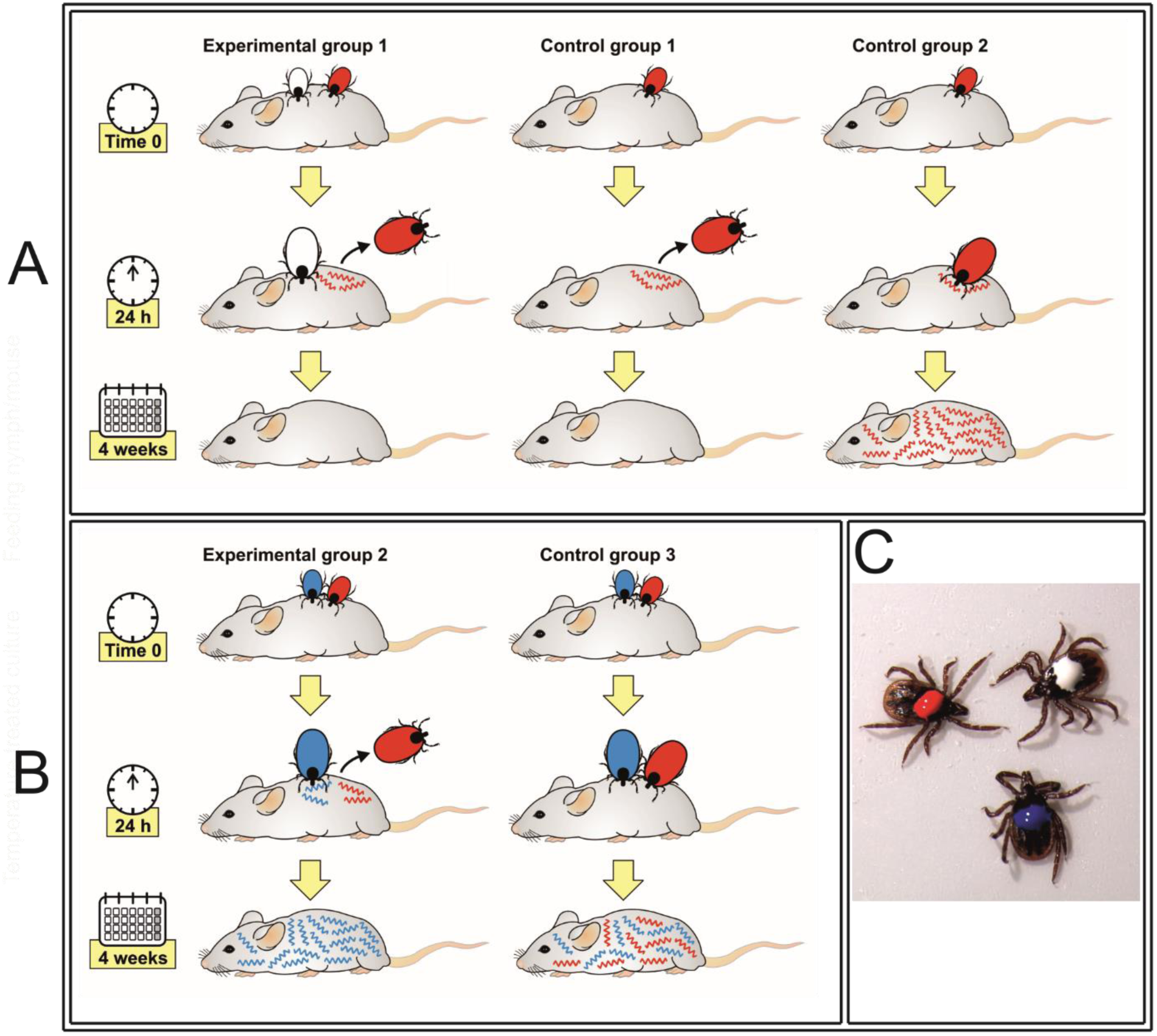
The role of tick saliva in *B. afzelii* survival. Presence of neither uninfected ticks (**A**) nor *B. burgdorferi* infected ticks (**B**) and their saliva is not sufficient for protection of early *B. afzelii* against their elimination by the host immune system.,(**C**) Differentially labeled *I. ricinus* nymphs. White: Uninfected nymph. Red: *B. afzelii* infected nymph. Blue: *B. burgdorferi* infected nymph.

A possible explanation of this unanticipated result might be that unlike the uninfected tick, the salivary glands of *Borrelia*-infected ticks express a different spectrum of molecules that assist their transmission and survival within the vertebrate host (14–16). Therefore, we also examined the protective effect of saliva from *Borrelia*-infected nymphs. The experimental setup was the same as above with one exception: In experimental group 2, nymphs infected with a different strain of *B. burgdorferi* were allowed to feed until repletion next to *B. afzelii*-infected nymphs that were removed after 24 hours. In control group 3, *B. afzelii*-infected and *B. burgdorferi*-infected nymphs were allowed to feed until repletion. Four weeks after repletion, mice were specifically examined for the presence of one or both *Borrelia* strains using *rrs–rrlA* IGS PCR amplification. All mice in experimental group 2 were positive for *B. burgdorferi*, while *B. afzelii* was not detected in any of analyzed murine tissues. All mice in control group 3 tested positive for both *B. afzelii* and *B. burgdorferi* (Fig. 6B). This result implies that the saliva from *B. burgdorferi* infected ticks was also not capable of ensuring survival of *B. afzelii* transmitted to mice at the early feeding stage.

### Infectivity by *B. afzelii* is gained in the midgut and changes during nymphal feeding

Another possible explanation for the delayed capability of *B. afzelii* to infect mice was that infectivity of the spirochetes changed during the course of nymphal feeding. To test the infectivity of *B. afzelii* during different phases of nymphal feeding, *B. afzelii* containing guts were dissected from unfed, 24 hour, 48 hour and 72 hour-fed *I. ricinus* nymphs and subsequently injected into C3H/HeN mice (5 guts/mouse). *B. afzelii* spirochetes from unfed nymphs were not infectious for mice. Spirochetes from nymphs fed for 24 hours infected 3 out of 5 inoculated mice and all mice became infected after the injection of spirochetes from 48 hour-fed nymphs. Interestingly, mice inoculated with spirochetes from nymphs fed for 72 hours established *B. afzelii* infection only in 1 out of 5 mice. This result suggests that the capability of *B. afzelii* spirochetes to infect mice is gained in the tick gut and peaks about the 2^nd^ day of feeding.

### Infectivity of *B. afzelii* is linked to differential gene expression during tick feeding and transmission

Previous research demonstrated that transmission of *B. burgdorferi* from *I. scapularis* to the host is associated with changes in expression of genes encoding outer surface proteins OspA, OspC or the fibronectin-binding protein BBK32 (7, 17–19). In order to examine whether the infectivity of *B. afzelii* may depend on expression of orthologous genes, we performed qPCR analysis to determine the status of *ospA*, *ospC* and *bbk32* expression by *B. afzelii* spirochetes in unfed and feeding *I. ricinus* nymphs as well as in murine tissues 4 weeks post infection. The gene encoding OspA was abundantly expressed in unfed ticks, down-regulated during tick feeding and was hardly detectable in mice. The *B. afzelii ospC* gene was lowly expressed in unfed *I. ricinus* nymphs. Its expression steadily increased during feeding, with the highest levels of *ospC* mRNA at the 3^rd^ day of feeding. Significant *ospC* expression was also detected in mice with a permanent *B. afzelii* infection. Similarly, a gradual up-regulation of *bbk*32 was evident with the progress of tick feeding and gene transcription was fully induced during mammalian infection (Fig. 7).

**FIG 7.**
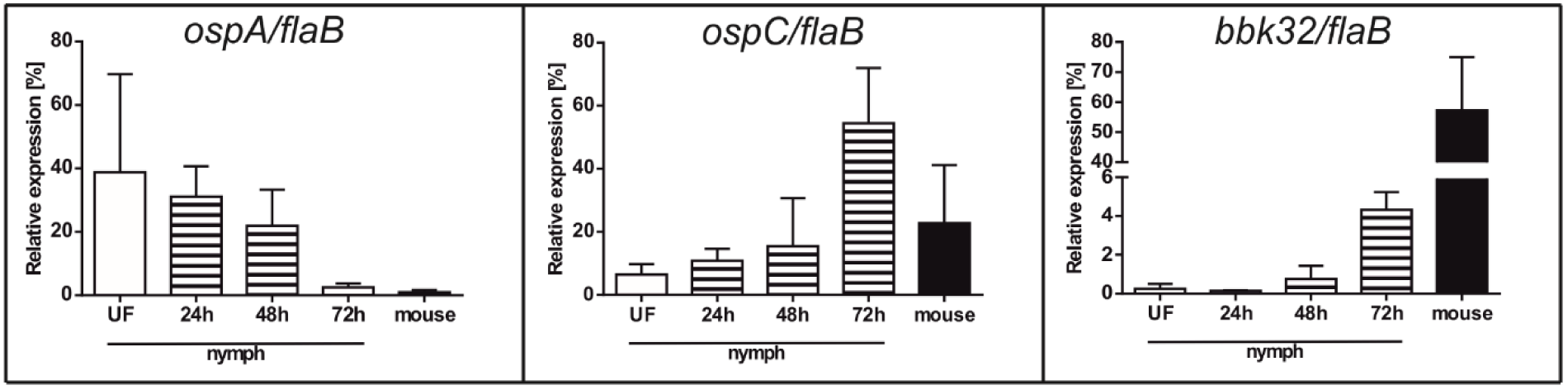
Comparative analysis of *ospA*, *ospC* and *bbk32* gene expression in *B. afzelii* spirochetes during tick feeding and mouse infection. Each datapoint represents the mean of 3 individually analyzed samples, and bars indicate standard errors of means.

## Discussion

Understanding the dynamics of *Borrelia* spirochete transmission is crucial for development of strategies for preventing Lyme disease. Recently, we managed to implement a reliable transmission model for European Lyme disease that involves the vector *I. ricinus* and the most common causative agent of borreliosis in Europe - *B. afzelii* spirochetes. This allowed us to quantitatively track the growth kinetics and infectivity of *B. afzelii* during the *I. ricinus* life cycle and compare it to data known for the *I. scapularis*/*B. burgdorferi*.

In nature, infection is acquired by larval or nymphal ticks feeding on an infected host. Absolute quantification of *B. afzelii* spirochetes during larval development and molting to nymphs revealed that *I. ricinus* larva imbibes relatively low spirochete numbers (~600 per tick). The number of *B. afzelii* then gradually increases during larval molting and reaches its maximum of about 20 000 spirochetes per tick two weeks after molting to nymphs. The level then stabilizes at about 10 000 spirochetes in starving nymphs (Fig. 2). This course of spirochetal burden is roughly in line with the data reported for *I. scapularis*/*B. burgdorferi* (6). However, compared to our observations, these authors described a dramatic decrease in *B. burgdorferi* numbers during *I. scapularis* molting. They speculate that it was due to depleted amounts of N-acetylglucosamine – an important building block of integumentary chitin but also a key component for spirochetal development. The limited availability of other nutrients might also be the reason for halted proliferation of spirochetes in molted nymphs. With its adoption of a parasitic lifestyle, the bacterium is an auxotroph for all amino acids, nucleotides and fatty acids. It also lacks genes encoding enzymes for the tricarboxylic acid cycle and oxidative phosphorylation (20, 21). Therefore, *Borrelia* spirochetes in the tick midgut are completely dependent on nutrients derived from ingested blood.

A striking difference between *I. ricinus*/*B. afzelii* and *I. scapularis*/*B. burgdorferi* was observed in spirochete numbers in the nymphal midgut during feeding. We found that *B. afzelii* numbers dramatically decrease from about ~10 000 spirochetes present in flat *I. ricinus* nymphs to only ~700 spirochetes in nymphs fed for three days (Fig. 3). This result is in sharp contrast with the data previously published for *I. scapularis*/*B. burgdorferi*. Using antibody-based detection, De Silva and Fikrig demonstrated that the total number of *B. burgdorferi* increased from several hundred in starved nymphs, to almost 170 000 spirochetes on the 3rd day of nymphal feeding (10). Later, these data were confirmed in a qPCR study showing that *B. burgdorferi* spirochetes in tick midguts increased six fold, from about 1 000 before attachment to about 6 000 at 48 hours after attachment (11).

It is commonly known that the risk of Lyme disease increases with the length of time a tick is attached. It was stated that *I. scapularis* ticks infected with *B. burgdorferi* removed during the first 2 days of attachment do not transmit the infection (11, 17). Our data show that *B. afzelii* spirochetes require less time to establish a permanent infection. Most mice became infected by 48 hours of attachment. This is in agreement with the previously published results showing that *B. afzelii* infected *I. ricinus* nymphs transmit the infection earlier than *B. burgdorferi* infected ticks (22).

Nevertheless, quantification by qPCR as well as microscopic examination of *B. afzelii* in the mouse dermis revealed that a substantial number of *B. afzelii* spirochetes enter the host earlier than they are able to develop a systemic infection (Fig. 5). This is in agreement with their significant decrease in the tick midgut during feeding (Figs. 3 and 4) and suggests that massive numbers of *B. afzelii* spirochetes leave the nymphs as early as the first day of feeding. The presence of spirochetes in mouse dermis prior to becoming infectious was also reported for *I. scapularis*/*B. burgdorferi.* Ohnishi et al. observed non-infectious spirochetes in skin samples from mice that were exposed to *B. burgdorferi* infected *I. scapularis* nymphs for less than 53 hours (17). Moreover, Hodzic et al. also reported the presence of *B. burgdorferi* spirochetes in four out of eight mice 24 hours after *I. scapularis* attachment (23). These data suggest that *Borrelia* spirochetes invade the host at very early time-points of tick feeding but early spirochetes are not able to develop a systemic infection. There could be two explanations for this observation. Firstly, bioactive molecules present in tick saliva are crucial for successful dissemination and survival of spirochetes within the host body. Therefore, the early spirochetes cannot colonize the host without sufficient protection and support of the tick saliva (24, 25). Secondly, early spirochetes that are transmitted to the vertebrate host are not infectious. A substantial body of work has been performed to elucidate the various tick bioactive molecules, mainly comprising a complex cocktail of salivary proteins that dampens the host’s defenses against blood loss and the development of inflammatory and complement reactions at the feeding site (26). Several tick molecules have been suggested to be crucial for *Borrelia* acquisition in ticks and transmission to the next host during subsequent feeding (reviewed in (27)). To test the role of tick saliva in survival of early spirochetes, we performed a co-feeding experiment in which the early *B. afzelii* spirochetes were under the protection of uninfected ticks or ticks infected with *B. burgdorferi* (Fig. 6). This experiment clearly showed that the presence of tick saliva is not sufficient for protection and survival of early spirochetes as all mice remained uninfected with *B. afzelii* spirochetes. Therefore, we tested how infectivity of *B. afzelii* changes during tick feeding. A number of studies provide solid evidence that *Borrelia* spirochetes change expression of their surface antigens during feeding and transmission to the host, making it possible for spirochetes to specifically adapt to the tick or the host environment, as required (7, 28, 29). Changes in gene expression of our model spirochete seem to be the main event that promotes increasing infectivity during tick feeding. *Borrelia afzelii* spirochetes in unfed ticks showed high levels of expression of *ospA* and negligible expression of *ospC* and *bbk32*. In this tick mode, spirochetes were not infectious for mice. As feeding progressed, *ospA* was downregulated and *ospC* and *bbk32* were upregulated, which correlated with increasing infectivity of *B. afzelii*.

The highest level of infection was observed in mice inoculated with spirochetes from 48 hour-fed nymphs. By this time, all mice had developed the infection. Interestingly, spirochetes from 72 hour-fed nymphs infected only one out of five mice. This decrease is likely associated with a concomitant, substantially reduced number of *B. afzelii* in the midguts of nymphs fed for 3 days (Figs. 3 and 4).

The route of *Borrelia* spirochete transmission has been the subject of controversy for a long time. In 1984, Burgdorfer suggested that spirochetal development in most ticks (*I. scapularis* and *I. ricinus*) occurs in the midgut. Additional tissues, including salivary glands, were considered to be free of spirochetes in most of the ticks. It was suggested that transmission occurs by regurgitation of infected gut contents or via saliva by ticks with a generalized infection (4). Benach et al. presented similar findings in their extensive histological study. They stated that *B. burgdorferi* are able to enter the hemocoel during the midfeeding period and develop a systemic infection in the hemolymph and central ganglion. However, *B. burgdorferi* were never seen within the lumen of the salivary gland or attached to cells of the salivary acini (2). Salivary route of Lyme disease transmission came into consideration since 1987, when Ribeiro and colleagues reported the presence of spirochetes in saliva of pilocarpine treated ticks (3) and then broadly accepted after microscopic detection of spirochetes within the salivary glands and ducts of fully fed *I. scapularis* nymphs (30). Nevertheless, the spirochete numbers present in salivary glands of *I. scapularis* nymphs are minuscule and hardly detectable (31, 32).

We have never observed *B. afzelii* present in salivary glands at any stage of tick feeding. The absence of spirochetes in salivary glands is surprising since large numbers of spirochetes were supposed to pass from the midgut to the feeding lesion during the three day course of nymphal feeding. Collectively, these results rather testify against the salivary transmission of *B. afzelii* by *I. ricinus*. Instead, we assume that regurgitation, or rather active reverse migration of motile spirochetes from midgut to the mouthpart (for which we coin the term ‘gut-to-mouth’ route) is the most probable way of transmission of *B. afzelii* from *I. ricinus* nymph to the host. This alternative way of *B. afzelii* transmission avoiding *I. ricinus* hemocoel and salivary glands is indirectly supported also by our recent research showing that silencing of tick immune molecules or elimination of phagocytosis in tick hemocoel by injection of latex beads had no obvious impact on *B. afzelii* transmission (33–35).

From our results we conclude that *B. afzelii* has a unique transmission cycle that, in many aspects, differs from the generally accepted salivary route of *B. burgdorferi* transmission by *I. scapularis*. *Borrelia afzelii* in flat *I. ricinus* nymphs represents a relatively abundant population of spirochetes. Once the tick finds a host, *B. afzelii* immediately start their transmission, most likely directly from the midgut to the feeding cavity. *B. afzelii* also seems to be less dependent on its tick vector. The main requirement for successful host colonization is the change in outer surface protein expression that occurs in the tick gut during the course of feeding. Spirochetes ‘switched’ to the proper, vertebrate mode are then able to survive within the host even if the tick is not present. The 24-48 hour time window between tick attachment and transmission of infectious spirochetes is the critical period in the whole process. These findings collectively point to the *Borrelia*-tick midgut interface as the correct target in our endeavours to combat Lyme disease transmission.

## Materials and Methods

### Laboratory animals

*Ixodes ricinus* larvae and nymphs were obtained from the breeding facility of the Institute of Parasitology, Biology Centre, Czech Academy of Sciences. Ticks were maintained in wet chambers with a humidity of about 95%, temperature 24°C and day/night period set to 15/9 hours. To prepare both infected and uninfected *I. ricinus* nymphs, the larvae were fed on either infected or uninfected mice, allowed to molt to nymphs, and after 4–6 weeks were used for further experiments. Inbred, pathogen free C3H/HeN mice (The Jackson Laboratory, Bar Harbor, ME, USA), were used for the pathogen transmission experiments.

### Ethics Statement

All experimental animals were treated in accordance with the Animal Protection Law of the Czech Republic No. 246/1992 Sb., ethics approval No. 161/2011. The animal experimental protocol was approved by the Czech Academy of Sciences Animal Care & Use Committee (Protocol Permit Number: 102/2016).

### Infection of mice and ticks

Low passage strains of *B. afzelii* CB43 (13), and *B. burgdorferi* SLV-2(36) were grown in BSK-H medium (Sigma-Aldrich, St. Louis, MO, USA) at 33°C for 5–7 days. Six weeks old female C3H/HeN mice were infected by subcutaneous injections of 10^5^ spirochetes per mouse. The presence of spirochetes in ear biopsies was verified 3 weeks post injection by PCR. Clean *I. ricinus* larvae were fed on infected mice until repletion and allowed to molt. Nymphs were considered to be infected if >90% of them were PCR positive.

### Nucleic acid isolation and cDNA preparation

DNA was isolated from individual larvae, nymphs, as well as murine tissues using a NucleoSpin Tissue Kit (Macherey-Nagel, Düren, Germany) according to the manufacturer’s protocol.

Total RNA was extracted from nymphs and murine tissues using a NucleoSpin RNA Kit (Macherey-Nagel) according to the manufacturer’s protocol. Isolated RNA (1 μg) served as a template for reverse-transcription into cDNA using Transcriptor High Fidelity cDNA Synthesis Kit (Roche, Basel, Switzerland). All cDNA preparations were prepared in biological triplicates.

### PCR

Detection of spirochetes in ticks, as well as in murine tissues was performed by nested PCR amplification of a 222 bp fragment of a 23S rRNA gene (37). PCR reaction contained 12.5 μl of FastStart PCR Master (Roche), 10 pmol of each primer, template (4 μl of DNA for the first round, 1 μl aliquot of the first PCR product in the second round), and PCR water up to 25 μl. Primers and annealing temperatures are listed in Table 1.

**Table 1.**
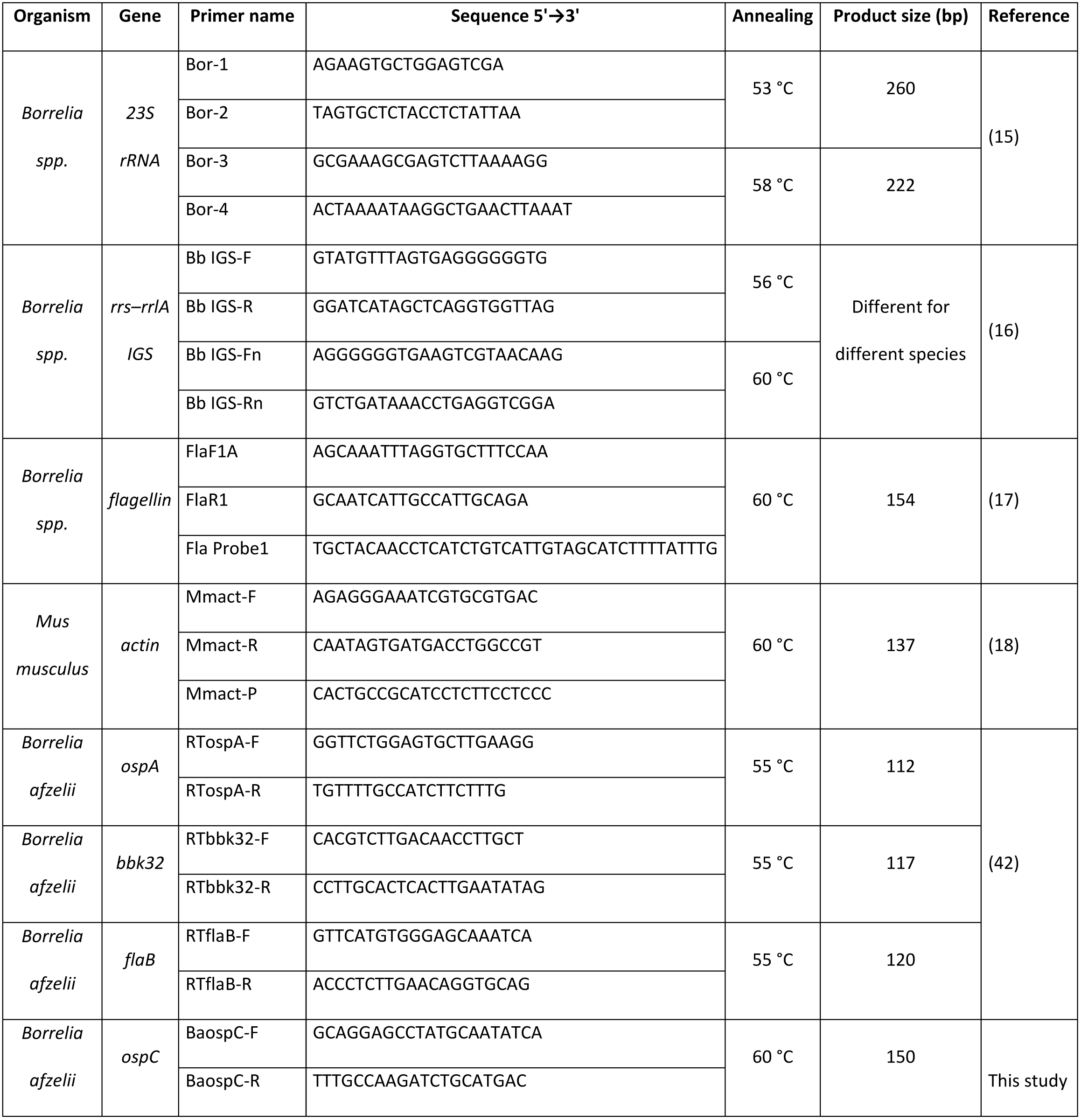
Primers and probes used in this study.

Differentiation of *B. afzelii* and *B. burgdorferi* strains was performed by nested PCR amplifying a part of *rrs–rrlA* IGS region (38). Reaction conditions were the same as above, primers and annealing temperatures are listed in Table 1.

### Quantitative real-time PCR

Total spirochete load was determined in murine and tick DNA samples by quantitative real-time PCR (qPCR) using a LightCycler 480 (Roche). qPCR reaction contained 12.5 μl of FastStart Universal Probe Master (Rox) (Roche), 10 pmol of primers FlaF1A and FlaR1, 5 pmol of TaqMan probe Fla Probe1 (39) (Table 1), 5 μl of DNA, and PCR water up to 25 μl. The amplification program consisted of denaturation at 95°C for 10 minutes, followed by 50 cycles of denaturation at 95°C for 15 sec and annealing + elongation at 60°C for 1 min.

Quantification of murine *β-actin* was performed using MmAct-F and MmAct-R primers and a MmAct-P TaqMan probe (40) (Table 1). Reaction and amplification conditions were the same as above. The spirochete burden in murine tissues was expressed as the number of spirochetes per 10^5^ of murine *β-actin* copies. The spirochete burden in ticks was calculated as the total number of spirochetes in the whole tick body.

cDNAs from *B. afzelii* infected *I. ricinus* nymphs as well as murine tissues served as templates for the quantitative expression analyses by relative qPCR. Reaction contained 12.5 μl of FastStart Universal SYBR Green Master, Rox (Roche), 10 pmol of each primer (Table 1), 5 μl of cDNA, and PCR water up to 25 μl. The amplification program consisted of denaturation at 95°C for 10 minutes, followed by 50 cycles of denaturation at 95 °C for 10 sec, annealing at 60 °C for 10 sec and elongation at 72°C for 10 sec. Relative expressions of *ospA*, *ospC* and *bbk32* were normalized to *flaB* using the ΔΔCt method (41).

### Preparation of murine and tick tissues for confocal microscopy

*Borrelia afzelii*-infected *I. ricinus* nymphs were fed on mice for 24 hours. Then, skin biopsies from the tick feeding site were dissected. Guts and salivary glands of unfed, 24 hour-fed, 48 hour-fed and fully fed nymphs infected with *B. afzelii* were dissected in phosphate buffer. Dissected tissues were immersed in 4% paraformaldehyde for 4 hours at room temperature. Tissues were then washed 3x20 min in PBS, permeabilized with 1% Triton X-100 (Tx) in PBS containing 1% BSA (Sigma) at 4°C, overnight. Next day, *Borrelia* spirochetes in tissues were stained with primary rabbit anti *B. burgdorferi* antibody (Thermo Fisher Scientific) 1:200 in PBS-Tx (0.1% Tx in PBS), for 4 hours at room temperature. Tissues were then washed 3x20 min in PBS-Tx and stained with Alexa Fluor 488 goat anti-rabbit, secondary antibody (Life Technologies, Camarillo, CA, USA), 1:500 in PBS-Tx, for 2 hours at room temperature. Tissues were counterstained with DAPI for 10 minutes and washed 2x10 min in PBS. Then, slides were mounted in DABCO and examined using an Olympus FluoView FV1000 confocal microscope (Olympus, Tokyo, Japan).

### Preparation of murine tissues for histology

*Borellia afzelii* infected or clean *I. ricinus* nymphs were fed on mice until repletion (5 mice/group, 10 nymphs/mouse). Four weeks later, murine tissues (skin, heart, urinary bladder) from *B. afzelii* infected and uninfected mice were fixed in 10% buffered formalin, embedded in paraffin using routine procedures. 3 μm thin sections were cut and stained with hematoxylin and eosin. Slides were examined using an Olympus BX40 light microscope (Olympus).

### Statistical analysis

Data were analyzed by GraphPad Prism 6 for Windows, version 6.04 and an unpaired Student’s t-test was used for evaluation of statistical significance. An Ap value of P<0.05 was considered to be statistically significant. Error bars in the graphs show the standard errors of the means.

## Acknowledgments

This work was primarily supported by the Czech Science Foundation grant No. 17-27393S to RS, and additionally by the grants 17-27386S, 18-01832S to OH and PK, European Union FP7 project Antidote (Grant Agreement number 602272), “Centre for research of pathogenicity and virulence of parasites” (No. CZ.02.1.01/0.0/0.0/16_019/0000759) funded by European Regional Development Fund (ERDF) and Ministry of Education, Youth and Sport, Czech Republic (MEYS). We acknowledge the excellent technical assistance of Jan Erhart, Zuzana Zemanová, and Adéla Palusová. Special thanks go to Prof. Jan Kopecký and Dr. Helena Langhansová who generously provided *B. afzelii* CB43 strain. We acknowledge Martina Hajdušková (www.biographix.cz) for the design of Fig. 6.

